# Pervasiveness of exoribonuclease-resistant RNAs in plant viruses suggests new roles for these conserved RNA structures

**DOI:** 10.1101/433672

**Authors:** Anna-Lena Steckelberg, Quentin Vicens, Jeffrey S. Kieft

**Affiliations:** Department of Biochemistry and Molecular Genetics, and RNA BioScience Initiative, University of Colorado Denver School of Medicine, Aurora, CO 80045, USA.

## Abstract

Exoribonuclease-resistant RNAs (xrRNAs) are discrete folded RNA elements that block the processive degradation of RNA by exoribonucleases. xrRNAs found in the 3′ untranslated regions (UTRs) of animal-infecting flaviviruses and in all three members of the plant-infecting *Dianthovirus* adopt a complex ring-like fold that blocks the exoribonuclease; this ability gives rise to viral non-coding subgenomic RNAs. The degree to which these folded RNA elements exist in other viruses and in diverse contexts has been unclear. Using computational tools and biochemical assays, we discovered that xrRNA elements are widely found in viruses belonging to the *Tombusviridae* and *Luteoviridae* families of plant-infecting RNA viruses, demonstrating their importance and widespread utility. Unexpectedly, many xrRNAs are located in intergenic regions rather than in the 3’UTR and some are associated with the 5′ ends of subgenomic RNAs with protein-coding potential, suggesting that xrRNAs with similar scaffolds are involved in the maturation or maintenance of diverse subgenomic RNAs, not just the ones generated from the 3′UTR.

## INTRODUCTION

During infection, positive-sense RNA viruses produce full-length genomic RNA and many produce subgenomic RNAs (sgRNA) that can encode viral proteins or act as “riboregulators” that interact with and influence the cellular and viral machinery to benefit viral infection^1–6^. Most viral sgRNAs are thought to be produced directly through transcription; however, recent discoveries show that some noncoding viral sgRNAs result from discrete RNA elements that block the progression of 5’ to 3’exoribonucleases (Figure 1)^7–15^. Discrete, compactly-folded exoribonuclease-resistant RNA (xrRNA) elements were first identified in mosquito-borne flaviviruses (e.g. Dengue virus, Zika virus, West Nile virus), where they protect the genome’s 3’ untranslated region (UTR) from degradation^8^. The resultant decay intermediates accumulate and comprise biologically active viral non-coding sgRNAs (Figure 1) 8, 9, 12, 16–21.

**Figure 1.**
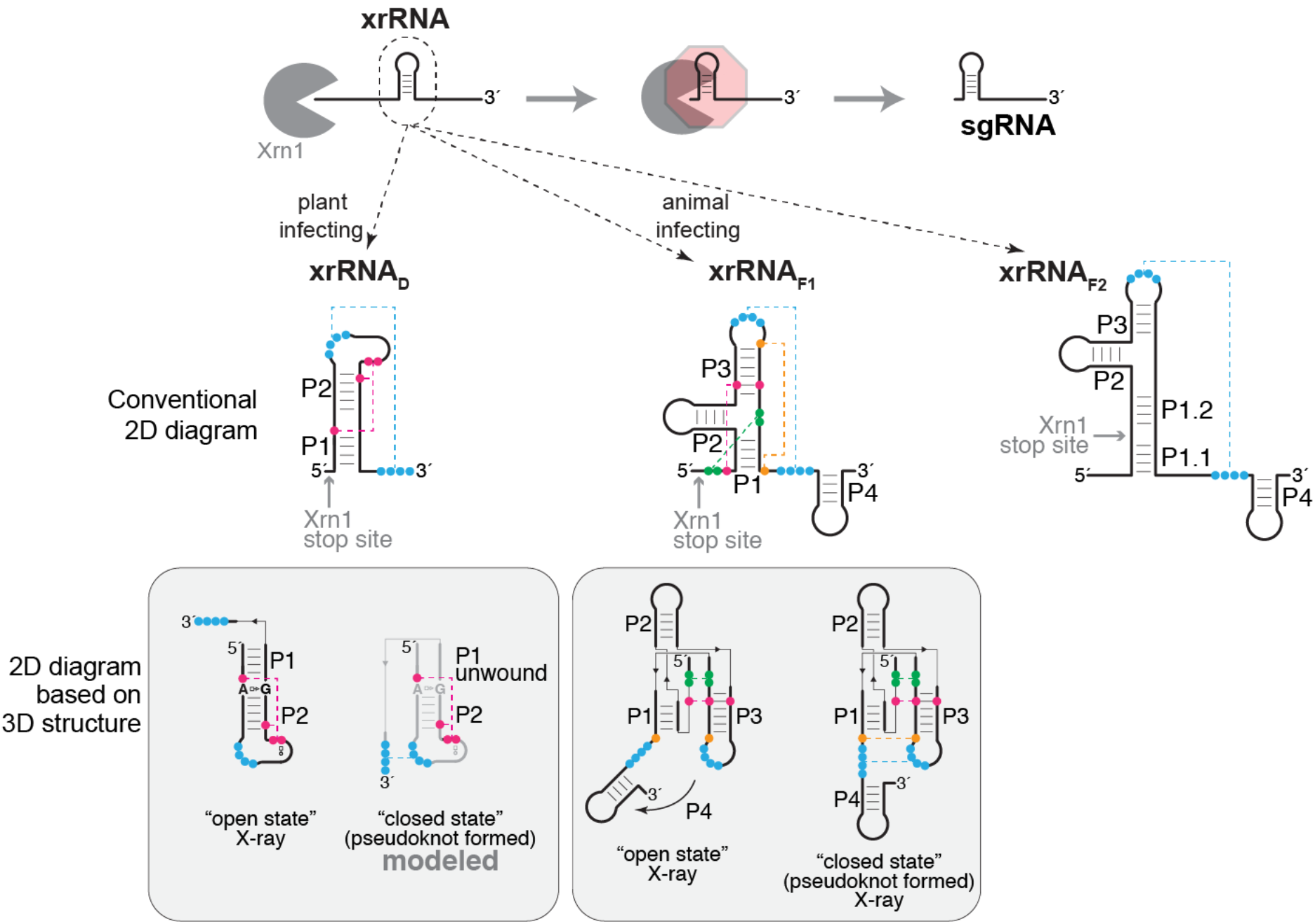
An expanding repertoire of structured RNAs for blocking exoribonuclease degradation. Top: xrRNAs adopt a three-dimensional structure that blocks the progression of 5′ to 3′ exoribonucleases such as Xrn1 (grey). In the case of flaviviruses and dianthoviruses, xrRNAs are in the 3′UTR and this results in accumulating sgRNAs that comprise the 3′UTR. Middle: Secondary structure diagrams are shown for the two classes of xrRNAs from flaviviruses (xrRNA_F1_ and xrRNA_F2_)^15, 24, 25^, and from dianthoviruses (xrRNA_D_)^26^. Secondary structure features are labeled, and nucleotides involved in tertiary interactions are shown in colors connected by dashed lines (pseudoknot shown in blue). Experimentally determined Xrn1 stop sites are indicated. Bottom: The grey shaded boxes below each secondary structure contain diagrams reflecting the currently available three-dimensional structures^22, 23, 26^. The A8-G33 pair is highlighted in the open state of the Sweet clover necrotic mosaic virus (SCNMV) xrRNA.

Extensive functional and high-resolution structural studies show that mosquito-borne flaviviral xrRNA (xrRNA_F_) function is conferred by a specific three-dimensional fold containing an interwoven pseudoknot stabilized by extensive conserved secondary and tertiary interactions; this creates an unusual ring-like conformation that protectively wraps around the 5’ end of the RNA structure^22, 23^. xrRNAs are found broadly within flaviviruses including those that are tick-borne, are specific to insects, or have no known vector^15, 24, 25^. Comparison of diverse xrRNA_F_ sequences revealed two classes; class I (xrRNA_F1_) is exemplified by mosquito-borne flaviviruses while class II (xrRNA_F2_) is found in diverse flaviviruses^25^. Although aligned xrRNA_F2_ sequences show patterns, their three-dimensional structures are unknown, as are the structures of recently reported xrRNAs from most other viral clades^7, 13^.

Recently, we structurally and functionally characterized xrRNAs from the 3′UTRs of dianthoviruses, plant-infecting positive-sense RNA viruses in the *Tombusviridae* family; similar to the xrRNA_F_, they function to produce a non-coding RNA derived from the viral 3′UTR^10,26^. Dianthoviral xrRNAs (xrRNA_D_) also rely on a pseudoknot that forms a protective ring-like structure ^26^, but they have very different sequences and secondary structures compared to xrRNA_F1_ and the ring is formed by a different set of interactions (Figure 1). Although xrRNA_F_ elements pervade the flaviviruses with associated sequence and structural diversity, xrRNA_D_ have only been identified in the three closely related members of the *Dianthovirus* genus. This raises the question of whether xrRNAs similar to xrRNA_D_ are more widespread and diverse than currently known, and thus if they may be an underappreciated way to produce or protect viral RNAs. Moreover, the only available xrRNA_D_ crystal structure is in an “open” conformation that likely represents a necessary folding intermediate before the pseudoknot forms ^26^ (Figure 1). Thus, we still do not know the full repertoire of secondary and tertiary interactions required to form and stabilize the exoribonuclease-resistant pseudoknot state of xrRNA_D_. The lack of diverse xrRNA_D_ sequences prevents conclusions about the role, prevalence, and structural diversity of this fold.

To begin to address these questions, we used a bioinformatic approach to identify more xrRNA_D_ sequences among plant viruses, identifying over 40 putative new xrRNA_D_-like elements in viruses belonging to the *Tombusviridae* and *Luteoviridae* families. *In vitro* assays show that these elements are indeed resistant to Xrn1 and analysis of these new xrRNAs reveals both conservation and variability. Furthermore, the genomic location of these new xrRNAs suggests new roles in the generation of sgRNA species that have protein-coding potential, providing evidence that xrRNA-based RNA maturation pathways may be more widespread than previously anticipated.

## RESULTS AND DISCUSSION

To search for xrRNA_D_-like elements, we used the *Infernal* software (Eddy lab), which enables screening of massive datasets of DNA sequences for conserved structure patterns with poor sequence conservation^27^. Because the *Dianthovirus* genus only comprises three members (Red clover necrotic mosaic virus (RCNMV), Sweet clover necrotic mosaic virus (SCNMV), and Carnation ringspot virus (CRSV))^26^, we expanded our search to other plant-infecting positive-sense RNA viruses. The initial search within a library of viral reference genomes (see Methods) identified two potential sequences among *Luteoviridae; Poleroviruses*: wheat leaf yellowing-associated virus isolate JN-U3 (GenBank ID # NC_035451; Infernal E-value = 0.00025, score = 44.3) and sugarcane yellow leaf virus (GenBank #NC_000874; Infernal E-value = 6.5, score = 24.2). With these sequences added to the alignment, subsequent searches identified > 40 candidates within the public repository of all available sequences for *Tombusviridae* and *Luteoviridae*, demonstrating how powerful this tool is for computationally identifying functional elements in viral RNAs^28^.

Alignment of the putative xrRNA_D_ –like elements evealed that their predicted secondary structures contain conserved helices P1, P2, and the pseudoknot, which are supported by covariation but have little sequence conservation (R-scape^29^ E-values for the 12 covarying base pairs in the stems and the pseudoknot are within 3.10^-4^–8.10^-13^ (95^th^ percentile = 1.10^-12^); Figure 2A). L1 and L2B are > 97% conserved in sequence. In the case of L1 and L2B, this is consistent with their role in creating a specific folded motif that promotes pseudoknot formation^26^. Also, two of the three nucleotides immediately upstream of the 3’ side of the pseudoknot are > 97% conserved, but their role is not obvious from the crystal structure of the open state. Likewise, the non-Watson-Crick A8-G33 base pair identified in the crystal structure (Figure 1) cannot be reconciled with the predominant presence of G at position 8 and G/A at position 33 in all the other sequences. These observations support the previous assertion that the crystallized open state represents a folding intermediate and that structural adjustments and additional interactions are present in the “closed” pseudoknot state.

**Figure 2.**
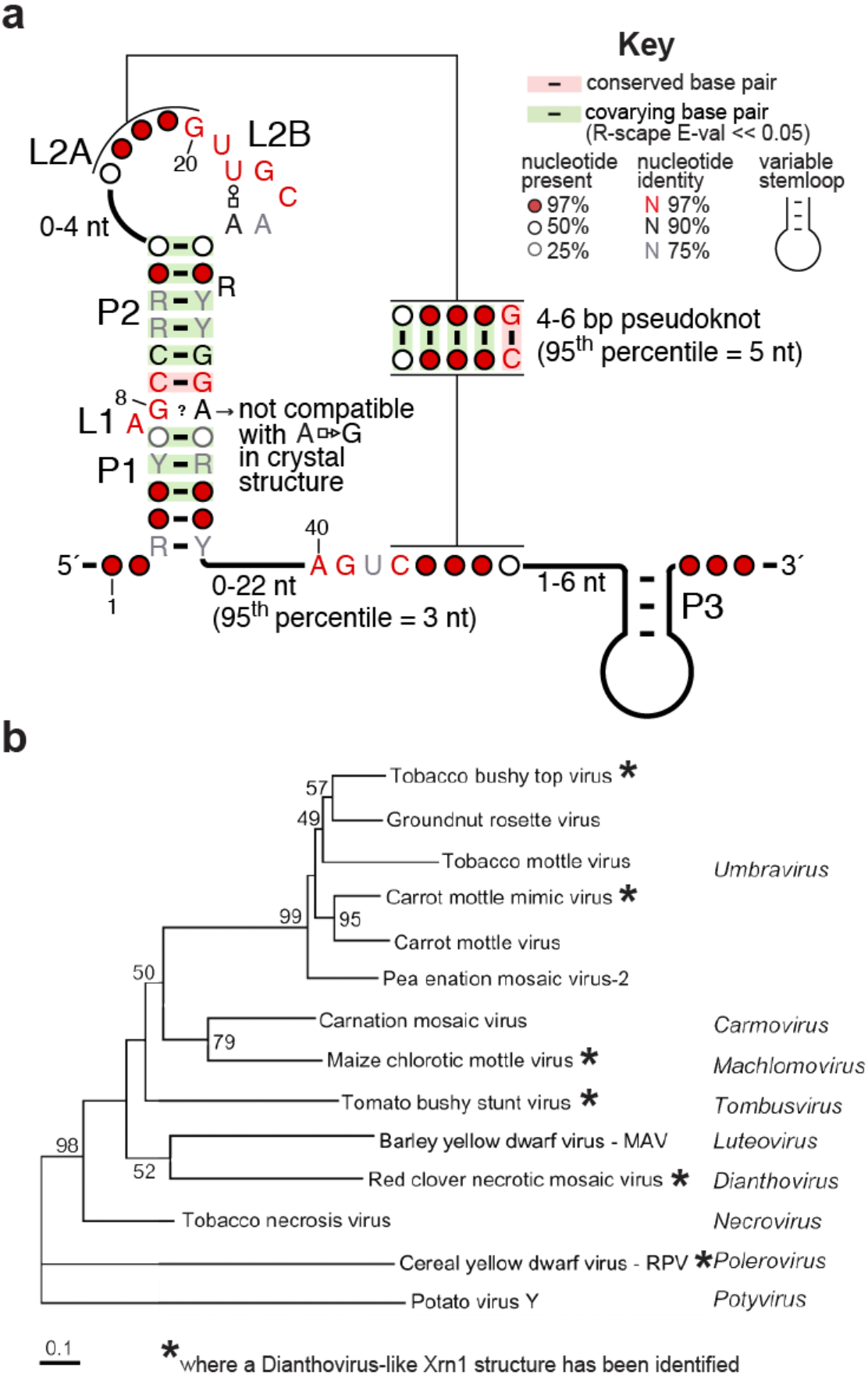
Widespread occurrence of Xrn1-resistant RNAs among plant viruses. (**a**) Consensus sequence and secondary structure of xrRNA_TL_ based on a comparative sequence alignment of 47 sequences of viruses belonging to the *Tombusviridae* and *Luteoviridae* families (shown in Figure S1). Y = pyrimidine; R = purine. Non-Watson-Crick base pairs are shown using the Leontis-Westhof nomenclature 43. The numbering is that of the crystal structure of the SCNMV xrRNA_D_ (now referred to as xrRNA_TL_)^26^. (**b**) Phylogenetic relationship between various plant viruses, based on the RNA-dependent RNA polymerase amino acid sequence^32^. The viruses and corresponding genera in which we identified xrRNA_TL_ structures are marked by a star. Numbers at the nodes refer to bootstrap values as percentages obtained from 2000 replications, shown only for branches supported by more than 40%. Branch length is proportional to the number of changes. Additional analysis will likely reveal xrRNA_TL_ elements in more of these viruses with additional sequence and structural variation.

Viruses with putative novel xrRNAs include members of the *Machlomovirus* and *Umbravirus* genera of the *Tombusviridae* family, as well as members of the *Polerovirus* and *Enamovirus* genera of the *Luteoviridae* family. To experimentally determine if these are authentic xrRNAs, we tested representative sequences from viruses of both families using our established *in vitro* Xrn1 resistance assay^11^. Specifically, *in vitro*-transcribed and purified RNA sequences from opium poppy mosaic virus (OPMV), Maize chlorotic mottle virus (MCMV), Potato leafroll virus (PLRV) Maize yellow dwarf virus-RMV (MYDV-RMV) and Hubei polero-like virus 1 (HuPLV1) were challenged with recombinant Xrn1. All RNAs stopped Xrn1 degradation similarly to RCNMV xrRNA_D_ (Figure 3A, B), demonstrating that they are authentic xrRNAs and do not require additional *trans*-acting proteins for function. Moreover, mutations to disrupt the putative pseudoknot in the MCMV, PLRV and HuPLV1 xrRNAs abolished Xrn1 resistance, while compensatory mutations that restore pseudoknot base pairing rescued the activity (Figure 3C-E). In addition, the mapped Xrn1 stop site is at the base of P1 in all newly identified xrRNAs, matching the xrRNA_D_ stop site (Figure 3F-H, Supplementary Figure S2). Overall, the conserved secondary structure (Figure 2B), the location of the exoribonuclease halt site, and the strict dependence on the pseudoknot for Xrn1 resistance suggest that these newly-identified xrRNAs use a similar molecular fold and mechanism as the xrRNA_D,_ thus we classify them as such, and hereafter refer to the class as xrRNA_TL_ (for *Tombusviridae* and *Luteoviridae*).

**Figure 3.**
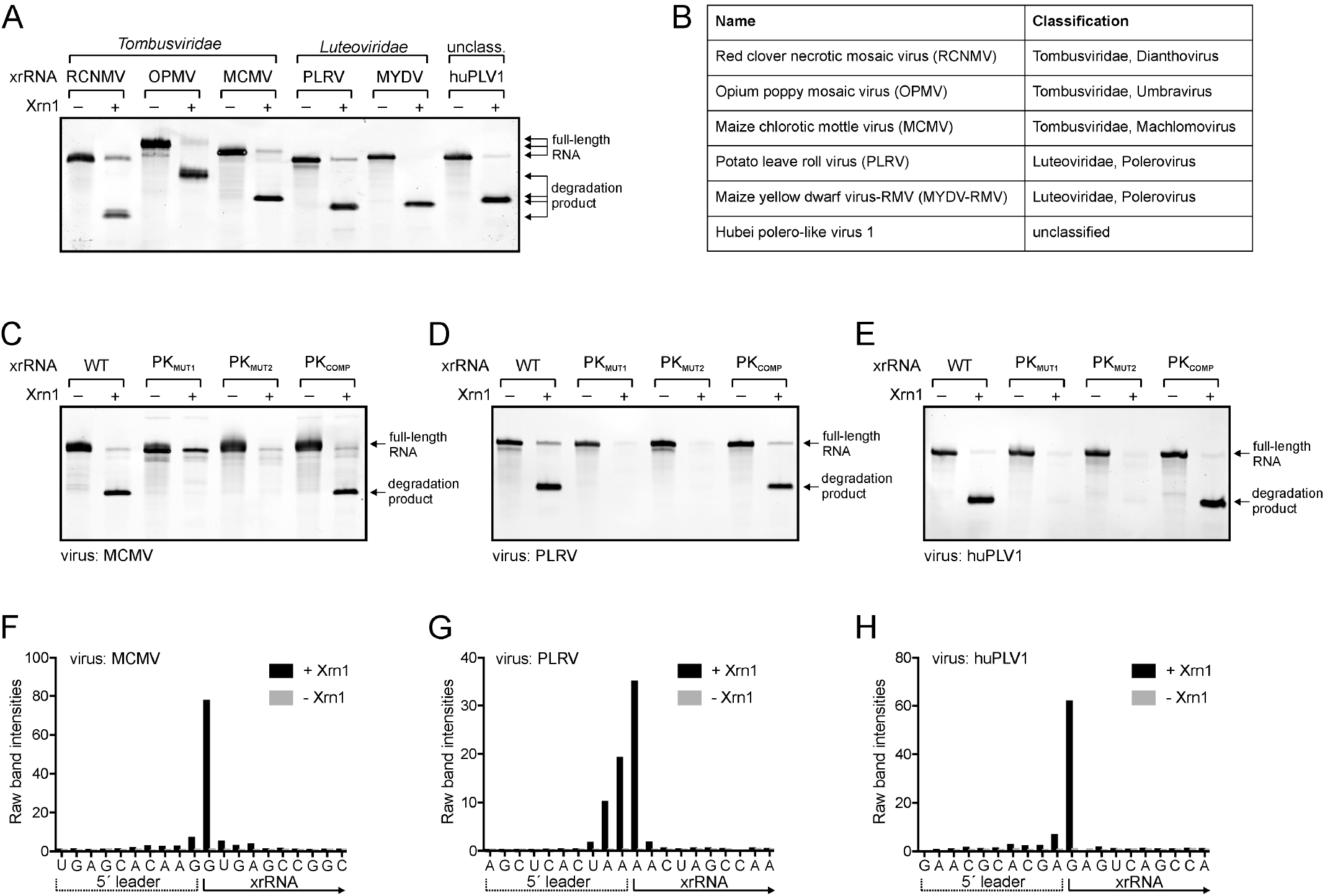
Biochemical characterization of representative plant virus xrRNA_TL_ elements. **(A)** *In vitro* Xrn1 resistance assay of putative xrRNA_TL_ from various plant RNA viruses (Table 1). The xrRNA from RCNMV was included as a positive control. Arrows indicate the size of full-length RNA and Xrn1-resistant degradation product. **(B)** Classification of viruses used in A (Table 1). **(C-E)** *In vitro* Xrn1 resistance assay of WT and PK mutant versions of MCMV (C), PLRV (D) and HuPLV1 (E) xrRNAs. **(F-H)** Reverse transcription (RT) mapping of the Xrn1 halt site. Distribution of RT products of Xrn1-resistant fragments of MCMV (F), PLRV (G) and HuPLV1 (H) degradation fragments. Experimentally validated halt sites are indicated on the secondary structure diagram for all tested xrRNA_TL_ in Figure S2.

A notable structural difference between diverse xrRNA_TL_ elements is that a subset of xrRNAs found in the *Tombusviridae* family (RCNMV, SCNMV, CRSV, OPMV, MCMV) possess a P3 stem-loop immediately downstream of the pseudoknot (Figure 2A; Tables 1 and S1). We previously showed that this part of the sequence is not required for Xrn1 resistance by xrRNA_TL_ *in vitro*^26^. Consistent with this, an analogous stem-loop (P4) found in xrRNA_F1_ is also dispensable *in vitro*; the crystal structure indicates it may stabilize the pseudoknot through stacking interactions (Figure 1)^22^. Thus, in xrRNA_TL_ coaxial stacking of P3 on P1/P2 could help to stabilize the RNA structure in the cellular context during infection.

**Table 1.** Selected set of plant viruses possessing an xrRNA_TL_. Viruses are grouped by their genomic context (last column). The complete list of sequences used for comparative sequence alignment is shown in Table S1.

| Name | Abbreviation | Classification | GenBank ID | Total ssRNA length (nt) | Genomic location* | Genomic context |
| --- | --- | --- | --- | --- | --- | --- |
| Red clover necrotic mosaic virus | RCNMV | Tombusviridae; Dianthovirus | NC_003756 | 3890 | 3461-3504 | 3' UTR |
| Sweet clover necrotic mosaic virus | SCNMV | Tombusviridae; Dianthovirus | NC_003806 | 3876 | 3446-3489 | 3' UTR |
| Maize chlorotic mottle virus (isolate KS1) | MCMV | Tombusviridae; Machlomovirus | NC_003627 | 4437 | 4101-4143 | 3' UTR |
| Opium poppy mosaic virus (isolate PHEL5235) | OPMV | Tombusviridae; Umbravirus | NC_027710 | 4230 | 3585-3629 | 3' UTR |
| Carrot mottle mimic umbravirus | CMoMV | Tombusviridae; Umbravirus | NC_001726 | 4201 | 2664-2706 | 74 nt to AUG from ORF3 |
| Chickpea chlorotic stunt virus | CpCSV | Luteoviridae; Polerovirus | NC_008249 | 5900 | 3489-3534 | 11 nt to AUA from ORF3a;<br>129 nt to AUG from ORF3-5 |
| Cowpea polerovirus 1 (isolate BE167) | CpPV1 | Luteoviridae; Polerovirus | NC_034246 | 5845 | 3380-3425 | 11 nt to CUG from ORF3a;<br>129 nt to AUG from ORF3-5 |
| Cotton leafroll dwarf virus | CoLRDV | Luteoviridae; Polerovirus | NC_014545 | 5866 | 3451-3499 | 13 nt to CUG from ORF3a;<br>131 nt to AUG from ORF3-5 |
| Cereal yellow dwarf virus-RPV | CYDV-RPV | Luteoviridae; Polerovirus | NC_004751 | 5723 | 3566-3622 | 14 nt to AUU from ORF3a;<br>132 nt to AUG from ORF3-5 |
| Maize yellow dwarf virus-RMV (Formerly BYDV) | MYDV-RMV | Luteoviridae; Polerovirus | NC_021484 | 5612 | 3335-3384 | 14 nt to ACG from ORF3a;<br>132 nt to AUG from ORF3-5 |
| Potato leafroll virus | PLRV | Luteoviridae; Polerovirus | NC_001747 | 5987 | 3509-3557 | 18 nt to AUA from ORF3a;<br>136 nt to AUG from ORF3-5 |
| Hubei polero-like virus 2 (strain QTM26674) | HuPLV2 | Unclassified | NC_033229 | 6083 | 3706-3753 | 133 nt to AUG from ORF3-5 |
| Beet western yellows luteovirus (strain bwv-1, isolate 28a) | BWVY | Luteoviridae; Polerovirus | L39983 | 973 | 341-389 | 135 nt to AUG from ORF3-5 |
| Hubei polero-like virus 1 (strain WHCC118254) | HuPLV1 | Unclassified | NC_032224 | 4213 | 3357-3410 | 135 nt to AUG from ORF3-5 |
| Sugarcane yellow leaf virus | ScYLV | Luteoviridae; Polerovirus | NC_000874 | 5899 | 3467-3512 | 136 nt to AUG from ORF3-5 |
| Beet western yellows virus | BWVY | Luteoviridae; Polerovirus | NC_004756 | 5666 | 3346-3393 | 138 nt to AUG from ORF3-5 |

\*xrRNA boundaries defined as the 1<sup>st</sup> nucleotide of the P1 stem and the last nucleotide of the pseudoknot.

The location of the new xrRNAs reveals unexpected variation (Figure 4). Only two of the newly identified xrRNAs are in the 3′UTR of the viral genome (Table 1), as are the previously characterized dianthoviral xrRNA_TL_ and xrRNA_F1-F2_. In MCMV, the first nucleotide of the P1 helix matches the 5’ end of sgRNA2^30^, thus this new xrRNA element probably blocks Xrn1 to generate non-coding sgRNAs derived from the 3′UTR, as with the dianthoviruses, flaviviruses, and other xrRNAs. However, for some members of the *Tombusviridae* family as well as for *Poleroviruses*, xrRNA_TL_ is located in an intergenic region, within 5–20 nt from the translation start site of ORF3a, and ∼ 135 nt from the start site of a readthrough protein encoded by ORF3–5 (our data suggest that ORF3a has not been annotated for all *Poleroviruses*; Table S1). ORF3a codes for protein P3a, which is essential for long-distance movement of the virus in plants^31^. Translation of ORF3a occurs from sgRNA1, generally at a non-AUG codon (Tables 1 and S1)^31–33^. This implies that these xrRNAs, rather than functioning in non-coding RNA production, act to produce or maintain protein coding RNAs (Figure 4); sgRNAs could be produced from full-length genomic RNAs without requiring a subgenomic promoter, or sgRNAs produced by other means could be protected from 5′ to 3′ degradation. Since the *Tombusviridae* and *Luteoviridae* families use 3′ proximal cap-independent translation enhancers (3′-CITEs) to initiate translation, uncapped sgRNAs with xrRNAs on their 5′ ends could still be translationally active^34,35^. Thus, these xrRNAs could be part of complex translation regulation mechanisms involving these 3′-CITEs and different sgRNAs^36^.

**Figure 4.**
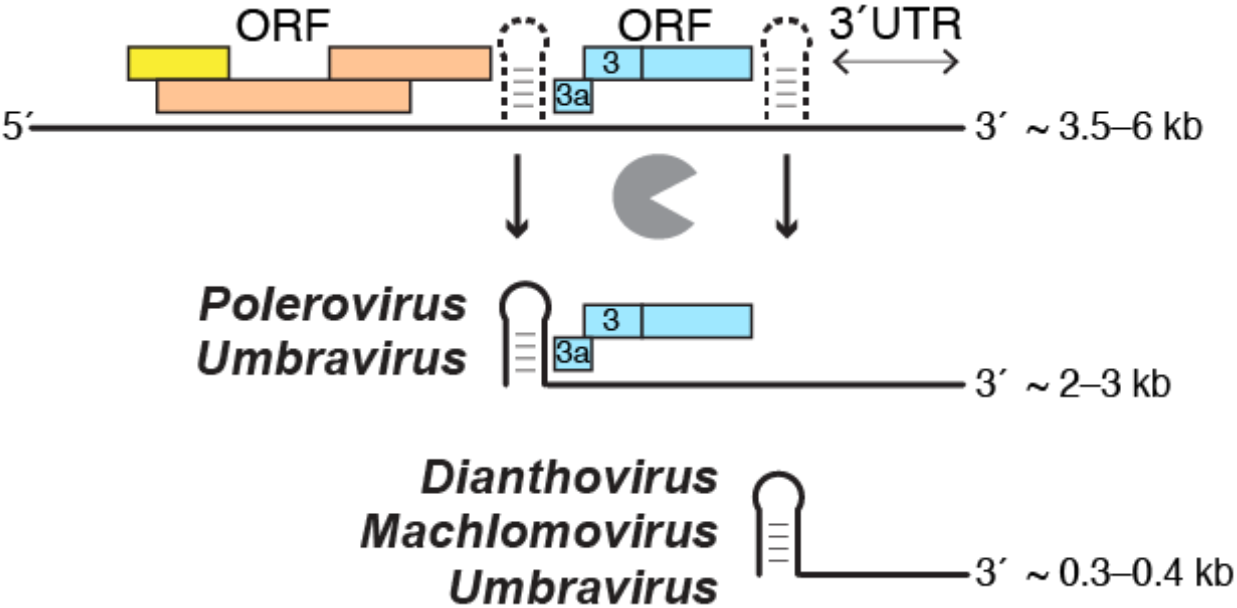
xrRNA_TL_ can produce or protect both coding and noncoding sgRNAs. The presence of xrRNA_TL_ in difference contexts suggests an expanded role for these elements. Shown here, full-length viral genomic RNA (top) could be processed by exonucleases that stop at xrRNAs (depicted as dashed structures) to yield both sgRNAs with protein coding potential (middle) and noncoding sgRNAs (bottom). Also, sgRNAs produced by subgenomic promoters could be “trimmed” or protected by xrRNAs (not shown). Note that only some *Umbraviruses* (e.g. OPMV) possess two xrRNA_TL_ elements. Colored boxes symbolize ORF organization in the plant viruses examined in this study.

Various roles for xrRNAs are possible, depending on their genetic context. The presence of xrRNAs in diverse locations within viral genomes suggests that new xrRNA scaffolds may emerge from analyzing sgRNA 5′ termini from other viruses, as certainly not all xrRNA elements were identified by the algorithm used here^5, 7, 37^ Intriguing candidates for novel xrRNA identification are viruses in which no obvious promoter elements for sgRNA production were identified, or viruses in which putative promoter sequences are downstream of the sgRNA 5′ end^1, 5, 30, 37^. Many questions remain that pertain to understanding the structural/sequence requirements for Xrn1 resistance, the degree to which structural variation is tolerated, and how sequence diversity is integrated into similar folds^38^. The now-expanded set of xrRNA_TL_ candidates provides a broader phylogeny for future bioinformatic and structural studies that will address these points.

## MATERIALS & METHODS

### Computational search

The published alignment with a total of three sequences (RCNMV, SCNMV and CRSV)^26^ was manually expanded in Ugene v. 1.29.0^39^ with two RCNMV variants (GenBank ID # J04357 and #AB034916) retrieved from a standard Nucleotide Blast search for “somewhat dissimilar sequences”(https://blast.ncbi.nlm.nih.gov/Blast.cgi?PAGE_TYPE=BlastSearch). Sequences were aligned to the conserved 3D-based secondary structure, omitting the pseudoknot, and exported in Stockholm format. This alignment was as follows:

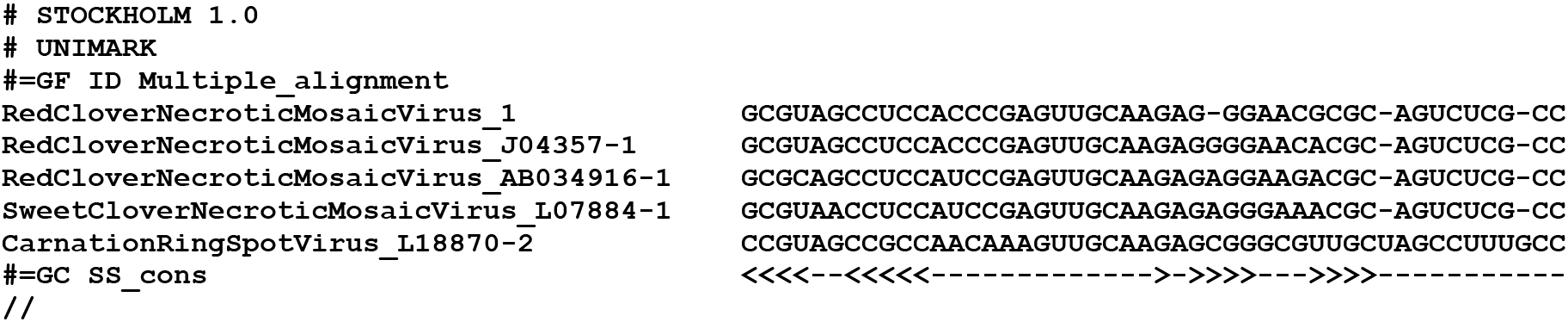

Using *Infernal* v. 1.1.2^27^ with default parameters, we searched for domains with similar structures and sequences within the complete reference genomes of viruses available from RefSeq, the NCBI Reference Sequence Database (https://www.ncbi.nlm.nih.gov/refseq/; downloaded on January 10, 2018). For subsequent iterations with Infernal, we searched the complete database of *Tombusviridae* and *Luteoviridae* available at GenBank (downloaded on July 3, 2018), using the following alignment:

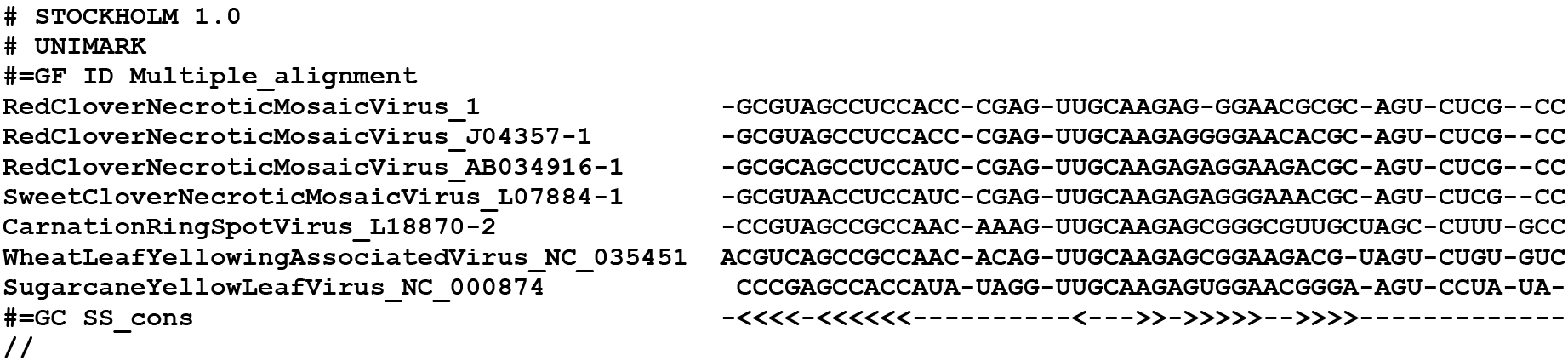

In Ugene, we systematically added new hits from *Infernal* to the alignment, only when they met the following criteria: (1) the sequence shows variation in more than 3–5 locations from the sequences already in the alignment; (2) the *Infernal* E-value is < 0.05; (3) the *Infernal* score is > 10; (4) the genomic context is coherent with that of the sequences already in the alignment. But a key in expanding the alignment further was to also analyze potential hits with a higher E-value / a lower score, as they would often correspond to positive hits but with a larger sequence or structure variation. By the time the alignment reached a size of 10–12 sequences, we were able to retrieve most of the sequences that made it into the final alignment through further iterations of *Infernal* searches and manual addition to the alignment. Hits for unclassified viruses were also retrieved from large-scale transcriptomics data of invertebrate and vertebrate-associated RNA viruses, using the deposited sequences^40,41^.

A statistical validation of the final proposed alignment of 47 sequences was performed using the latest version of R-scape available at http://eddylab.org/R-scape/^29^ (last accessed on August 17, 2018). The corresponding conserved structure and sequence patterns were rendered using R2R v. 1.0.5^42^.

### Design of RNAs for *in vitro* assays

DNA templates for *in vitro* transcription were gBlocks ordered from IDT, cloned into pUC19 and verified by sequencing. RNA constructs for Xrn1 degradation assays contained the xrRNA sequence plus ∼30 nucleotides of the endogenous upstream sequence (‘leader sequence’) to allow loading of the exoribonucleases. Below are the sequences used in *in vitro* Xrn1 degradation assays with the T7 promoter underlined, the leader sequence in italic and the first protected nucleotides (experimentally validated as described below) in bold. Lower case letters indicate extra nucleotides inserted to allow better transcription.

### OPMV xrRNA

<u>TAATACGACTCACTATA</u>*GGAATTGCCTCCACCAGTAACTAAACCCA****A*C**CACAGCCAAGCATTAA GTTGCAAGCGTTGGAGTGGCAGGCTTAACGTCCGACAGTACGACAACTGCGG

### MCMV xrRNA

<u>TAATACGACTCACTATA</u>*GGTTCCAGGCCCAGGGCTGGCAAATCATTGAGCACAAG***G**TGAGCCG GCATGAGGTTGCAAGACCGGAACAACCAGTCCTTCTGGCAGAGTCCTGCCAA

### PLRV xrRNA

<u>TAATACGACTCACTATAg</u>*GCCACCACAAAAGAACACTGAAGGAGCTCACTA***AA**ACTAGCCAAGC ATACACGAGTTGCAAGCATTGGAAGTTCAAGCCTCGT

### MYDV-RMV xrRNA

<u>TAATACGACTCACTATAg</u>*GTCCAGAAACAAAAAGTTTAAAACAG****A*A***GCTCTCA*AGTCAGCCAGGC AAATTCGAGTTGCAAGCACTGGATGACCTAGTCTCGATA

### HuPLV1 xrRNA

<u>TAATACGACTCACTATAg</u>*GCCACAAAACGAATAAAGGAAGAACGCACGA***G**AGTCAGCCAAACA AACACAAGTTGCAAGTGTTGGAGACTCATTCTAGTCTTGT

### RNA preparation

DNA templates for *in vitro* transcription were amplified by PCR using custom DNA primers (IDT) and Phusion Hot Start polymerase (New England BioLabs). 2.5 mL transcription reactions were assembled using 1000 µL PCR reactions as template (∼0.2 µM template DNA), 6 mM each NTP, 60 mM MgCl2, 30 mM Tris pH 8.0, 10 mM DTT, 0.1% spermidine, 0.1% Triton X-100, T7 RNA polymerase and 2 µL RNasin RNase inhibitor (Promega) and incubated overnight at 37°C. After inorganic pyrophosphates were precipitated by centrifugation, the reactions were ethanol precipitated and purified on a 7 M urea 8% denaturing polyacrylamide gel. RNAs of the correct size were gel-excised, eluted overnight at 4°C into ∼40 mL of diethylpyrocarbonate (DEPC)-treated milli-Q filtered water (Millipore) and concentrated using Amicon Ultra spin concentrators (Millipore). Mutations were introduced using mutagenized custom DNA reverse primers.

### Primers used in this study (mutated residues in bold)

OPMV_WT_rev 5′-CCGCAGTTGTCGTACTGTCGG-3′

OPMV_PKmut1_rev 5′-CCGCAGTTGTCGTACTGTCGGACG**AATT**GCCTGCCACTCCAACGC-3′

OPMV_PKmut2_rev 5′-CCGCAGTTGTCGTACTGTCGGACGTTAAGCCTGCCACTCCAACGCTTGCAAC**AATT**TGCTT GGCTGTGGTTGG-3′

OPMV_PKcomp_rev 5′-CCGCAGTTGTCGTACTGTCGGACG**AATT**GCCTGCCACTCCAACGCTTGCAAC**AATT**TGCTT GGCT GTGGTTGG-3′

MCMV_WT_rev 5′-TGGCAGGACTCTGCCAGAAGG-3′

MCMV_PKmut1_rev 5′-TGGCAGGACTCTGCCAG**CTCC**ACTGGTTGTTCCGGTCTTGC-3′

MCMV_PKmut2_rev 5′-TGGCAGGACTCTGCCAGAAGGACTGGTTGTTCCGGTCTTGCAA**GGAG**ATGCCGGCTCACC TTGTGCTC-3′

MCMV_PKcomp_rev 5′-TGGCAGGACTCTGCCAG**CTCC**ACTGGTTGTTCCGGTCTTGCAA**GGAG**ATGCCGGCTCACC TTGTGCTC-3′

PLRV_WT_rev 5′-ACGAGGCTTGAACTTCCAATGC-3′

PLRV_PKmut1_rev 5′-**TGCT**GGCTTGAACTTCCAATGCTTGC-3′

PLRV_PKmut2_rev 5′-ACGAGGCTTGAACTTCCAATGCTTGCAAC**AGCA**GTATGCTTGGCTAGTTTTAGTG-3′

PLRV_PKcomp_rev 5′-**TGCT**GGCTTGAACTTCCAATGCTTGCAAC**AGCA**GTATGCTTGGCTAGTTTTAGTG-3′ MYDV-RMV_WT_rev 5′-TATCGAGACTAGGTCATCCAGTGC-3′

huPLV_WT_rev 5′-ACAAGACTAGAATGAGTCTCC-3′ huPLV_PKmut1_rev 5′- **TGTT**GACTAGAATGAGTCTCCAACACTTGC-3′

huPLV_PKmut2_rev 5′- ACAAGACTAGAATGAGTCTCCAACACTTGCAAC**AACA**GTTTGTTTGGCTGACTCTCG-3′

huPLV_PKcomp_rev 5′- **TGTT**GACTAGAATGAGTCTCCAACACTTGCAAC**AACA**GTTTGTTTGGCTGACTCTCG-3′

### *In vitro* Xrn1 resistance assays

4 µg RNA was resuspended in 40 µL 100 mM NaCl, 10 mM MgCl2, 50 mM Tris pH 7.5, 1 mM DTT and re-folded at 90°C for 3 minutes then 20°C for 5 minutes. 3 µL recombinant RppH (0.5 µg/µL stock) was added and the samples were split into two 20 µL reactions (-/+ exoribonuclease). 1 µL of the recombinant Xrn1 (0.8 µg/µL stock) was added where indicated. All reactions were incubated for 2 hrs at 30°C using a thermocycler. The degradation reactions were resolved on a 7 M urea 8% denaturing polyacrylamide gel and stained with ethidium bromide.

### Mapping of the exoribonuclease halt site

To determine the Xrn1 stop site at single-nucleotide resolution, 30 µg *in vitro*-transcribed RNA was degraded using recombinant RppH and Xrn1 as described above (the reaction volume was scaled up to 300 µL, and 20 µL of each enzyme was used). The degradation reaction was resolved on a 7 M urea 8% polyacrylamide gel, then the Xrn1-resistant degradation product was cut from the gel and eluted overnight at 4°C into ∼20 mL of diethylpyrocarbonate (DEPC)-treated milli-Q filtered water (Millipore) and concentrated using Amicon Ultra spin concentrators (Millipore). Once recovered, the RNA was reverse-transcribed using Superscript III reverse transcriptase (Thermo) and a 6-FAM (6-fluorescein amidite)-labeled sequence-specific reverse primer (IDT) with an (A)20 –stretch at the 5′ end to allow cDNA purification with oligo(dT) beads. 10 µL RT reactions contained 1.2 pM RNA, 0.25 µL 0.25 µM FAM-labeled reverse primer, 1 µL 5x First-Strand buffer, 0.25 µL 0.1 M DTT, 0.4 µL 10 mM dNTP mix, 0.1 µL Superscript III reverse transcriptase (200 U/µL) and were incubated for 1 hour at 50°C. To hydrolyze the RNA template after reverse transcription, 5 µL of 0.4 M 4 NaOH was added and the reaction mix incubated at 90°C for 3 min, followed by cooling on ice for 3 min. The reaction was neutralized by adding 5 µL of acid quench mix (1.4 M NaCl, 0.57 M HCl, 1.3 M sodium acetate pH 5.2), then 1.5 µL oligo(dT) beads (Poly(A)Purist MAG Kit (Thermo)) were added and the cDNA purified on a magnetic stand according to the manufacturer’s instructions. The cDNA was eluted in 11 µL ROX-HiDi and analyzed on a 3500 Genetic Analyzer (Applied Biosystems) for capillary electrophoresis. A Sanger sequencing (ddNTP) ladder of the undigested RNA was analyzed alongside each degradation product as reference for band annotation.

## CONTRIBUTIONS

Q.V., A.-L.S., J.S.K. designed and analyzed research; Q.V. performed the computational search; A.-L.S. performed the biochemical experiments; Q.V., A.-L.S. & J.S.K wrote the paper.

## ACKNOWLEDGEMENTS

We gratefully acknowledge David Farrell for IT support; the Kieft lab members for useful discussions; and W. Allen Miller and David Costantino for careful reading of the manuscript.

